# Widespread remodelling of proteome solubility in response to different protein homeostasis stresses

**DOI:** 10.1101/692103

**Authors:** Xiaojing Sui, Douglas E. V. Pires, Shuai Nie, Giulia Vecchi, Michele Vendruscolo, David B. Ascher, Gavin E. Reid, Danny M. Hatters

## Abstract

The accumulation of protein deposits in neurodegenerative diseases involves the presence of a metastable subproteome vulnerable to aggregation. To investigate this subproteome and the mechanisms that regulates it, we measured the proteome solubility of the Neuro2a cell line under protein homeostasis stresses induced by Huntington Disease proteotoxicity; Hsp70, Hsp90, proteasome and ER-mediated folding inhibition; and oxidative stress. We found one-quarter of the proteome extensively changed solubility. Remarkably, almost all the increases in insolubility were counteracted by increases in solubility of other proteins. Each stress directed a highly specific pattern of change, which reflected the remodelling of protein complexes involved in adaptation to perturbation, most notably stress granule proteins, which responded differently to different stresses. These results indicate that the robustness of protein homeostasis relies on the absence of proteins highly vulnerable to aggregation and on large changes in aggregation state of regulatory mechanisms that restore protein solubility upon specific perturbations.

## INTRODUCTION

The protein homeostasis system is vital for cell function, as it ensures that proteins are properly translated, folded, trafficked to their correct cellular locations, and eventually degraded in a tightly controlled and timely manner. As a major task for this system is to prevent damaged and misfolded proteins from accumulating, it has been hypothesized that, when this system becomes unbalanced, proteins can become prone to misfolding leading to their mislocalisation and accumulation as aggregates (1–3). Dysfunctional protein aggregation and protein homeostasis imbalance are central pathological features of common neurodegenerative diseases including Alzheimer’s, Parkinson’s, Huntington’s, and motor neuron diseases (2, 4–7).

Different neurodegenerative diseases are characterised by the presence of signature proteins in the characteristic deposits formed in the brains of affected patients. These proteins include TDP-43 and FUS in motor neuron disease, tau and Aβ in Alzheimer’s disease, α-synuclein in Parkinson’s disease, and huntingtin (Htt) in Huntington’s disease. It has been suggested that these aggregation-prone proteins, which may be affected by mutations (e.g. Htt), post-translational modifications (e.g. tau phosphorylation), or environmental changes, lead to oversubscription of quality control resources that overloads the protein homeostasis system (8). This imbalance then triggers a cascade of protein folding defects that broadly impairs proteome function. In cell models expressing mutant Htt exon 1 (Httex1) for example, key chaperone systems are sequestered into inclusions formed by Httex1, which depletes the resources of the protein homeostasis system (9–11). Studies employing *C. elegans* models have shown that the aggregation of polyQ proteins can cause temperature-sensitive mutant proteins, which are metastable in their native states, to aggregate (10) which is consistent with a reduced capacity of the quality control system. Similarly, in ageing, different genetic backgrounds and environmental stresses can alter the efficiency of the protein homeostasis system (12, 13).

Here we sought to address three outstanding questions related to the factors capable of compromising protein homeostasis. The first question is how does aggregation of mutant Httex1, which previously has been suggested to unbalance protein homeostasis, impact the aggregation state of the wider proteome? The second is which proteins in the proteome are metastable to aggregation under different triggers of protein homeostasis stress? And the third is whether there is a subproteome that consistently requires a functional protein homeostasis machinery to remain folded and soluble, and if so, how is this subproteome regulated? Our results indicate that a substantial proportion of proteome undergoes large changes (upwards and downwards) in solubility in response to stress, but also that each stress is associated with an articulated stress response that affects a different part of the metastable subproteome. Our data suggests the majority of the changes arise from the functional remodelling of protein-ligand complexes in adaptation (or response) to the stress, and that the changes are highly specific to the different stress factors.

## RESULTS

### Httex1 mutation, and subsequent aggregation, distinctly remodels proteome solubility

To investigate how protein homeostasis imbalance alters the aggregation state of the proteome we employed a neuronal-like cell model system (mouse Neuro2a cells) and a quantitative proteomic workflow inspired by the work of Wallace et al. (14). In essence, the approach involved a fractionation strategy based on centrifugation of cell lysates prepared using a mild non-ionic detergent based lysis conditions (IGEPAL CA-630), with subsequent quantitative proteome analysis to monitor changes in the abundances of individual proteins between the supernatant versus the pellet, resulting from each stress (**Fig 1**). Quantitation was performed using a reductive dimethylation labelling approach with n=4 biological replicates and detection by nano reversed-phase liquid chromatography coupled with tandem mass spectrometry (MS/MS). We measured the changes in abundance of proteins in the total starting material (Experiment 1 in **Fig 1A**) and applied two reported methods to measure changes in solubility (14), which are anticipated to provide a different dynamic range of detection to proteins of different abundances and solubilities. Δ *pSup* was defined as the change in proportion of protein in the supernatant by subtracting the proportion of protein in the stress (*pSup_Stress_*) from the supernatant of control (*pSup_Control_*) (Experiments 2 and 3 in **Fig 1B**). We also measured the changes in the pellet fraction directly as the ratio of proteins in the stress:control treatments (Experiment 4 in **Fig 1B**). Hereon, we use the term “solubility” to indicate the changes in protein mass measured by this specific experimental framework.

**Figure 1.**
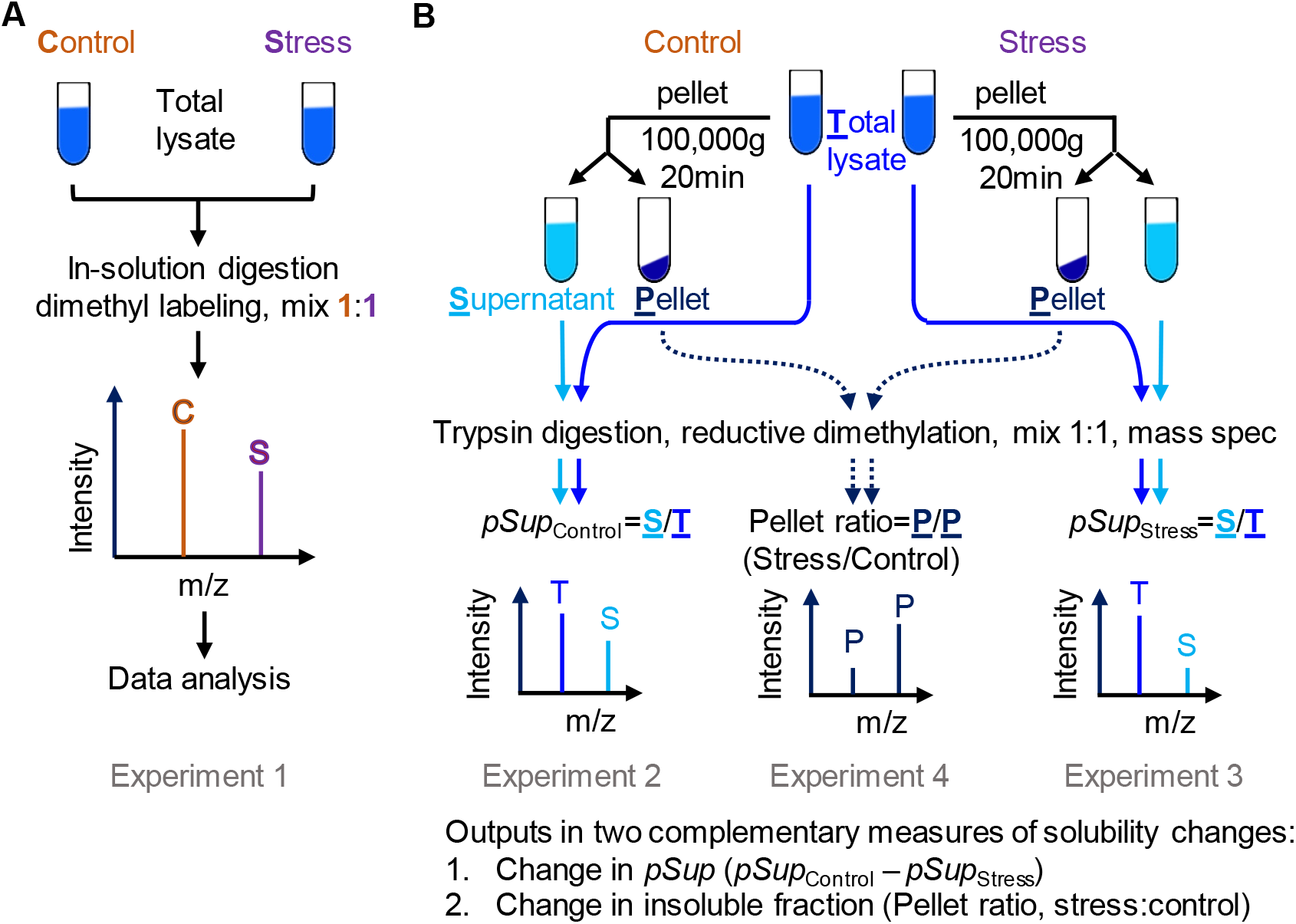
Proteomic workflow for quantitative measurements of stress-related proteome solubility changes. (**A**) Strategy to measure changes in total proteome abundances due to stress (Experiment 1). (**B**) Strategies to measure changes in solubility by a combination of xperiments (Experiments 2–4).

Our first protein homeostasis stress model examined the effect of Httex1 mutation and aggregation state on the overall proteome solubility. Huntington’s disease mutations lead to expansion of a polyglutamine (polyQ) sequence in Httex1 to lengths longer than 36Q, whereas the wild-type protein is typically less than 25Q (15). PolyQ-expansion causes Httex1 to become highly aggregation prone, which manifests as intracellular inclusion bodies as the disease progresses (16–18). We and others have used transient expression constructs of Httex1 fused to fluorescent proteins as models for replicating essential features of the disease, including protein homeostasis stress (19–23). Mutant Httex1-fluorescent protein constructs progressively form large cytosolic inclusions in cell culture over time.

We employed the flow cytometry method of Pulse Shape Analysis to purify cells expressing mutant Httex1-mCherry into those with inclusions (i) from those without inclusions (ni; dispersed uniformly in the cytosol) at matched median expression levels (21, 24). This strategy enabled us to assess how the aggregation state of mutant Httex1 (97Q ni and i) affected proteome solubility compared to a wild-type state (25Q ni – note that 25Q does not form aggregates) (**Fig 2A and Fig S1**).

**Figure 2.**
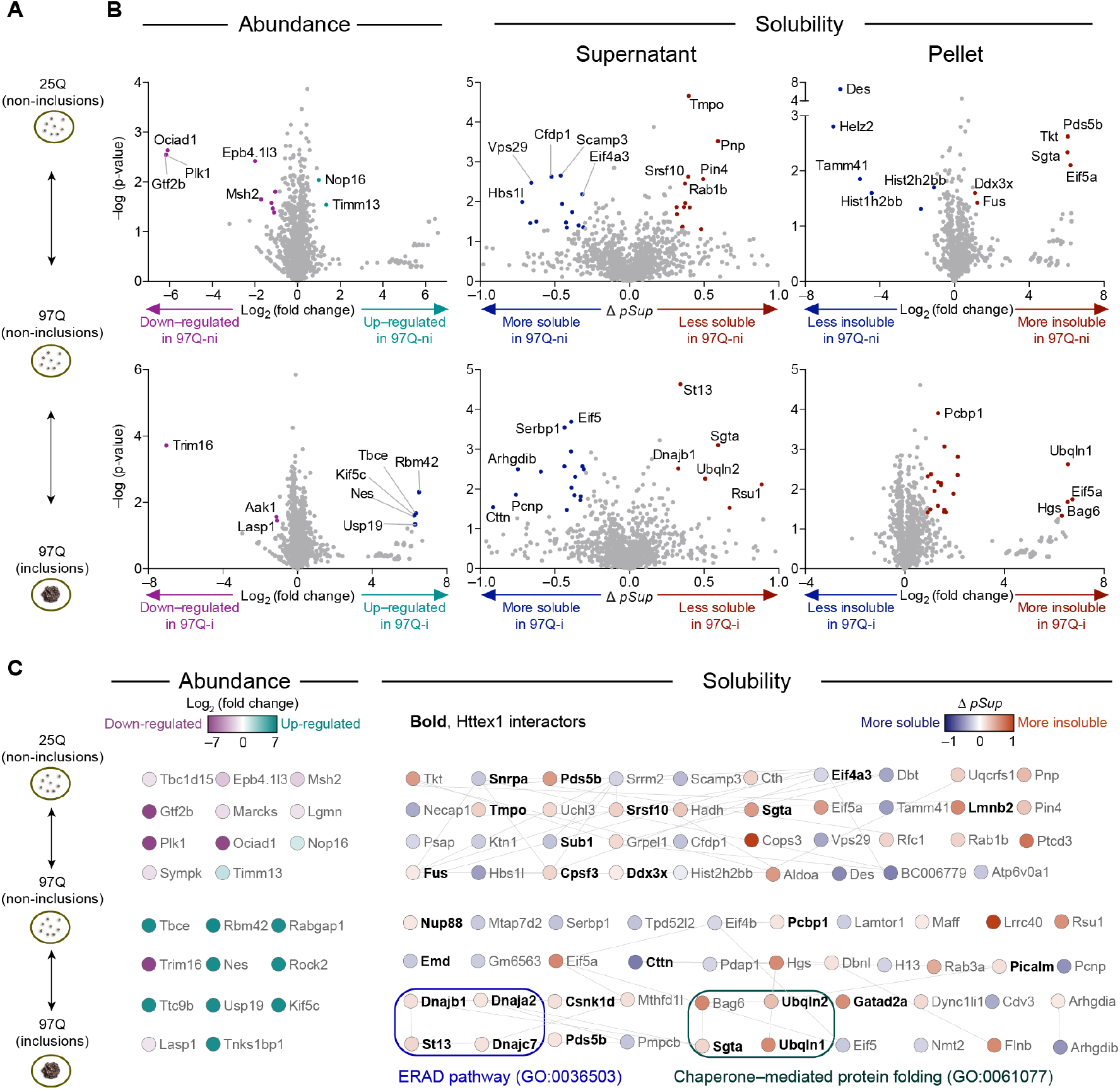
Impact of a Httex1 mutation and subsequent aggregation on solubility of endogenous proteins. (**A**) Neuro2a cells transiently expressing Httex1-GFP for wild-type polyQ (25Q) and mutant polyQ (97Q) were presorted into three groups by PulSA with matched expression levels as shown. (**B**) Volcano plots of protein abundances and solubility, aligned to schematic in panel A. Grey-colored points indicate proteins below the threshold of significance (2-fold change and P value of 0.05 for abundance and *Δ pSup* value of 0.3, Pellet ratio log_2_ value of 1 and P value of 0.05). (**C**) Protein networks represented by STRING (v.11) analysis with medium confidence interactions. Selected significantly enriched Gene Ontology terms are annotated. All enriched Gene Ontology terms are included in Supplementary Data 1

First, we assessed the protein abundances using the Experiment 1 pipeline from **Fig 1A**. A total of 2013 proteins were identified (**Table S1**). Only a handful of proteins (22) significantly changed abundance between the 25Q-ni, 97Q-ni and 97Q-i samples (**Fig 2B**). None of these had known protein-protein interactions (**Fig 2C**) and the dataset was enriched with only one Gene Ontology (GO) term (cytoskeleton GO:0005856, **Supplementary Data 1**). However, several of the proteins have reported roles in Huntington’s disease-relevant mechanisms. Further details of the functional details of these proteins are included in an expanded discussion in **Supplementary Note 1**.

For the assessment of protein solubility changes, we observed 2519 proteins^1^ (**Fig 2B**). For the comparison between 97Q-ni and the wild-type 25Q-ni we observed a slightly higher proportion of proteins that decreased solubility in 97Q-ni (17 more soluble and 22 less soluble). Likewise, a slightly higher proportion of proteins became more insoluble once Httex1 formed inclusions (97Q-ni versus 97Q-i) (16 more soluble and 25 less soluble).

Of the proteins that significantly changed solubility due to dispersed or inclusion-localized 97Q Httex1, almost one-third (24 out of 78) were previously reported as interactors of Httex1, 5 of which became more soluble (23) and 19 more insoluble (23, 25–28) (**Fig 2C**, *shown in bold*). One interesting feature was that 11 of the Httex1-interacting proteins are reported to localize to stress granules including Cttn, Pds5b, Cpsf3, Ddx3x, Dnajc7, Eif4a3, Ubqln2, Nup88, Pcbp1, Fus, and Srsf10 (29–34). 8 other stress granule proteins were also observed to change in solubility, including Helz2, Mthfd1, Serbp1, Eif5a, Eif4b, Cdv3, Pdap1, and Flnb. Of note is the broader role for stress granule abnormalities involved with misfolded proteins and neurodegenerative diseases (35–37). A GO analysis revealed 15 terms enriched in the proteins that changed their solubility when mutant Httex1 97Q formed inclusions. This set of terms included chaperone-mediated protein folding (GO:0061077) and ERAD pathway (GO:0036503) (full list of enriched GO terms in **Supplementary Data 1**). Collectively, these data suggested that mutant Httex1 causes two major effects on the proteome. The first is a substantial remodeling of stress granule structures both before and after aggregation into inclusions, which includes some elements becoming more soluble and some less soluble. The second is that the quality control systems involved in ER stress and protein misfolding appear to be selectively remodeled to become less soluble as Httex1 inclusions form, which is consistent with the inclusions recruiting chaperones and other quality control machinery in attempts to clear them (9, 23).

### Different triggers of protein homeostasis stress invoke distinct functional remodelings of proteome solubility

To investigate whether the proteins that changed solubility upon Httex1 aggregation are relevant to protein homeostasis stress more generally, we expanded our analysis to examine proteome solubility changes associated with 5 other triggers of protein homeostasis stress that have previously reported roles leading to protein misfolding and aggregation. These stresses included three specific inhibitors of key protein homeostasis hubs (Hsp70, Hsp90 and the proteasome) whereby defects are reported in models of Huntington’s disease, protein aggregation, degradation of misfolded proteins and-or other markers of protein homeostasis stress (9, 22, 23, 38–43) and two exogenous stress states that reflect pathology observed in neurodegenerative disease settings and protein aggregation in cell models (namely, oxidative stress and ER stress) (44–50).

The Hsp70 chaperone system was targeted by the small molecule inhibitor Ver-155008, which binds to the ATPase domain of Hsp70 family proteins (*K_d_* of 0.3 μM and *IC*_50_ of 0.5 to 2.6 μM) (51, 52). We have previously demonstrated Ver-155008 impairs protein homeostasis in cell culture models and can increase the aggregation propensity of an ectopically expressed metastable bait protein (38). Hsp90 was targeted with the ATP binding competitor novobiocin, which binds to the C-terminal nucleotide binding pocket to inhibit activity (40, 42, 43, 53). Unlike other drugs that target the N-terminal domain of Hsp90, novobiocin does not induce a heat shock response (54). We showed previously that novobiocin can unbalance the protein homeostasis system and induce the aggregation of a metastable bait protein (38). Proteasome activity was targeted with the inhibitor MG132 (55), which induces the formation of ubiquitin-positive aggregates in several cell lines (56–61). ER stress was triggered using the N-linked glycosylation inhibitor tunicamycin, which impairs flux of folding via the calnexin-calreticulin folding pipeline (62–64). Oxidative stress was induced with arsenite (65, 66).

The changes in protein abundance from these treatments are shown in **Fig S2A** (the full proteomics datasets are summarized in **Supplementary Tables 2–6**). The treatments led to changes consistent with their function based on GO assignment as well as protein-protein interactions (**Fig 3A;** complete list of GO terms in **Supplementary Data 1**). Of note was that many more proteins were observed to have changed solubility (upwards and downwards) than had changed abundance, which suggests that protein solubility change, rather than changes in protein expression, is a particularly substantial response to stress (Volcano plots in **Fig S2B**). Similar to what was observed for protein abundance changes, the GO terms indicated functional groupings anticipated from protein functions related to the treatment (**Fig 3B** for three examples of Hsp90 inhibition, proteasome inhibition and oxidative stress; **Fig S3** for the other two of Hsp70 inhibition and ER stress). Indeed, richer insights, particularly for oxidative stress and Hsp90 inhibition, was observed in the GO terms and details of the network of protein-protein interactions resulting from proteome solubility changes, than what was determined from the abundance changes, further supporting the conclusion that solubility is a particularly sensitive measure of the functional remodeling of protein complexes in response to stress (**Fig 3B**). For example, MG132 treatment indicated enrichment for Proteolysis (GO: 0006508) as anticipated. An effect on proteasome activity was also indicated by MG132 increasing the abundance of ubiquitin and proteasome subunits (**Fig S4A**). Almost all of the Proteolysis GO terms were associated with proteins becoming more insoluble, suggesting that the proteasome-degradation machinery forms larger molecular weight complexes when the proteasome is inhibited, which is consistent with the prior knowledge that proteasome inhibition induces the formation of ubiquitin- and proteasome-enriched cellular aggregates (67) (**Fig 3B**). Further insight into this functional remodeling of the proteasome machinery was provided by the substantial decrease in solubility of total cellular ubiquitin, even though an overall increase in total ubiquitin abundance was observed (**Fig S4B**).

**Figure 3.**
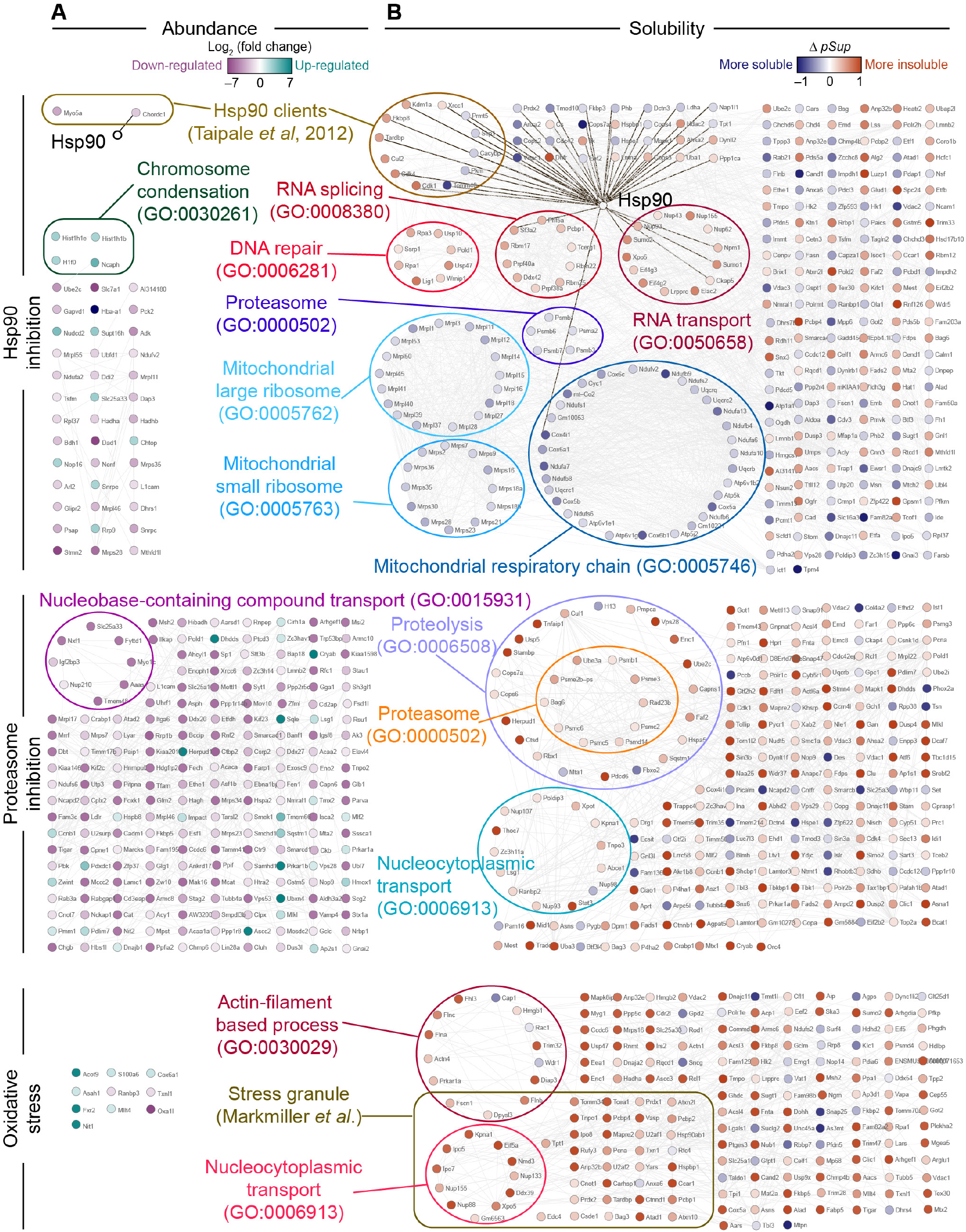
Impact of three protein homeostasis stresses on proteome abundance and solubility. Protein-protein interaction analysis of proteins with significant changes in abundance (**A**) or solubility (**B**) due to protein homeostasis stresses (Hsp90, proteasome inhibition and oxidative stress, top view) was performed built-in String (v.11) in Cytoscape (v3.7.1) at the medium confidence. Selected significantly enriched Gene Ontology (GO) terms are annotated. All enriched Gene Ontology terms are included in Supplementary Table S2. Hsp90 interactions were manually added based on String and shown with thicker black connectors. Stress granule curated list from (30).

Another notable finding from this analysis was the ability to extract novel information on the effect of Hsp90 inhibition on assembly states of macromolecular machines. Novobiocin did not change the levels of Hsp90 or proteins involved in the heat shock response, but decreased known Hsp90 client proteins, in accordance with previous studies (**Fig 3A**) (68, 69). However, GO and network analysis of the changes in solubility not only identified many more Hsp90 clients than those detected from expression level analysis, but also revealed large increases in solubility of proteins that form the proteasome, mitochondrial ribosome, DNA repair machinery, RNA splicing machinery, RNA transport machinery and respiratory chain complexes – details that are were less apparent from the analysis of the changes in proteome levels alone (**Fig 3B**). One of the major roles of Hsp90 is to mediate the assembly and stability of protein complexes (39) and in particular the 26S proteasome (70). Hence, our data suggests that the impairment of Hsp90 prevents subunits to properly assemble into large molecular weight machines that sediment under our experimental protocol, not only in the proteasome, but in the other major complexes of mitochondrial ribosome, respiratory chain complexes and potentially glycolysis-related enzymes and RNA and DNA-related processes. Of note is the mitochondrial respiratory chain, which contains five multimeric membrane-anchored complexes (71). We observed more than half of identified subunits of mitochondrial respiratory complex I, III and IV becoming more soluble after novobiocin treatment, suggesting a failure of these complexes to assemble into their mature state as part of large membrane-anchored complexes, which are anticipated to partition into the insoluble fraction under our pelleting regime.

### Consensus features of metastable subproteomes changing solubility under stress

To address the fundamental question of whether there is a common metastable subproteome that is more aggregation-prone under any condition of stress due to protein misfolding, we next sought to assess whether stress increased the net insolubility of the proteome by measuring the protein mass concentration in the supernatants before versus after pelleting. None of the stresses yielded a net change in the overall solubility (assessed by t-test of individual treatments)^2^. However, there was a significant difference (mean decrease of 3.8%) when the stresses were compared to controls in a paired one-tailed t-test (p = 0.03), with the statistical test justified on the hypothesis that stress increases insolubility of the proteome. These results suggest that aggregation arising from misfolded proteins increases marginally under stress, but does not reflect a dramatic accumulation of misfolded protein states that aggregate (**Fig 4A and Fig S5**).

**Figure 4.**
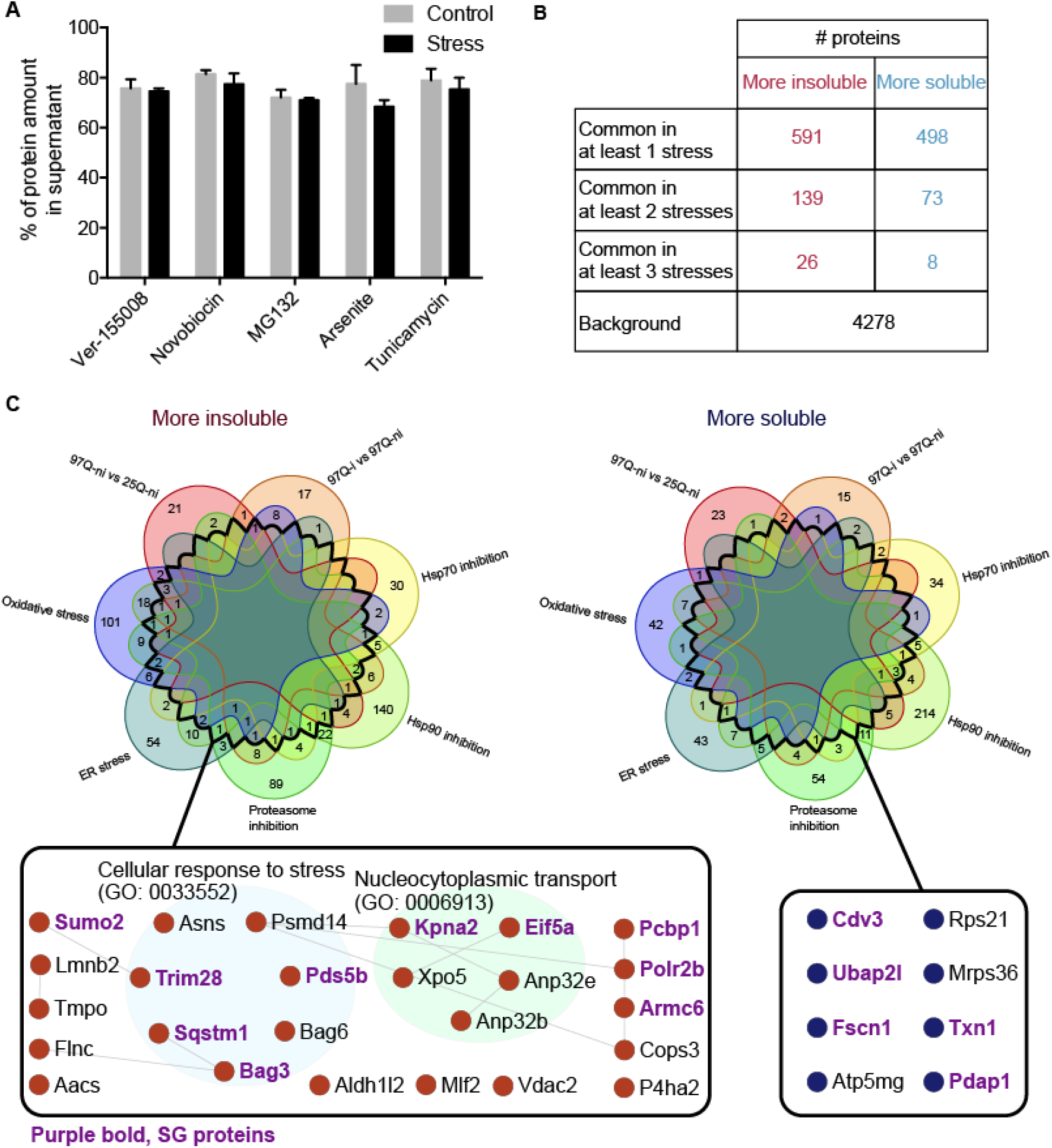
Dynamic remodeling of proteome solubility involving a core enrichment of proteins involved in nucleocytoplasmic transport and stress granules. (**A**) Overall proteome solubility changes due to treatments. Comparison of the proportion of protein amount in the supernatant fraction out of the total lysate measured by BCA assay between the control and treatment groups. Error bars represent SD. N= 3 or 4. Statistical significance was calculated by one-tailed paired t-test. P = 0.0297. (**B**) List of proteins found in common as shown. (**C**) Venn diagram presents the overlaps of more insoluble or soluble proteins across 6 stresses. The area enclosed by thick black line represents the overlap regions with at least 3 stresses. Protein-protein interaction of common proteins that became more insoluble (red) and more soluble (blue) in at least 3 stresses are shown. Purple bold represents known SG proteins.

At the individual protein level, there were no proteins that consistently decreased or increased in solubility across all 6 stresses using our criteria for a significant change. Of the proteins detected in all experiments, 591 proteins significantly decreased solubility by any one of the stress treatments, which represents 13.8% of the proteins detected (4278). In addition, 498 proteins had significantly increased solubility (11.6%). Altogether, the proteins that changed solubility was over one quarter of the proteome (25.4%) with a significant bias to proteins becoming more insoluble (Fishers Exact test p = 0.0028). Collectively these data indicate that significant remodeling in proteome solubility occurs under stress, but that most of the increase in insoluble protein load is counterbalanced by other proteins increasing in solubility.

Despite this widespread change in solubility, the changes were largely specific to the type of stress invoked. For example, we observed that only 26 proteins became more insoluble in at least 3 stresses, which represents just 0.6% of the proteome detected (**Fig 4B**) while 139 proteins (2.6% of the proteins) became more insoluble in at least 2 stresses. Only three proteins, Pcbp1, Bag3, Sqstm, were more insoluble in 4 stresses, and only one protein, asparagine synthetase (Asns), was more insoluble in 5 of the stresses (**Supplementary Table 7**). Likewise, for proteins that became more soluble, we observed only a low number of proteins (8) that became more soluble in at least 3 of the stresses (0.2% of the proteome detected) (**Fig 4B and Supplementary Table 7**). No protein was found to be more soluble in more than 3 stresses. Expanding the analysis to 2 stresses yielded 73 proteins (1.7% of the proteome).

GO analysis of the proteins that changed solubility in 2 or more stresses illuminated the key mechanisms involved (**Fig 5A**). The most enriched molecular function terms for the more insoluble protein list were proteasome-activating ATPase activity (82.6-fold enrichment), proteasome binding (36.7-fold), structural constituent of nuclear pore (35.9-fold), ATPase activator activity, (9.9-fold), heat shock protein binding (9.8-fold) ubiquitin protein ligase binding (6.0-fold). These functions are squarely consistent with protein quality control mechanisms being engaged to respond to proteome folding stress. Other major biological process and cellular component terms of enrichment include lamin filament (99.0-fold), nuclear pore outer ring (66-fold), mRNA export from nucleus (22.0-fold) and protein import into nucleus (14.6-fold), which are consistent with previously reported findings that protein folding stress more generally impacts nucleocytoplasmic transport mechanisms (32, 72–77). For the proteins that become more soluble, we observed terms highly enriched for mitochondrial activity including ATPase activity, coupled to transmembrane movement of ions, rotational mechanism (46.9-fold), proton-exporting ATPase activity (40.0-fold), proton transmembrane transporter activity (15.5-fold), ATP biosynthetic process (21.8-fold), mitochondrial protein complex (9.7-fold). These are consistent with mitochondrial complexes failing to assemble upon inhibition of Hsp90 (**Fig 3**).

**Figure 5.**
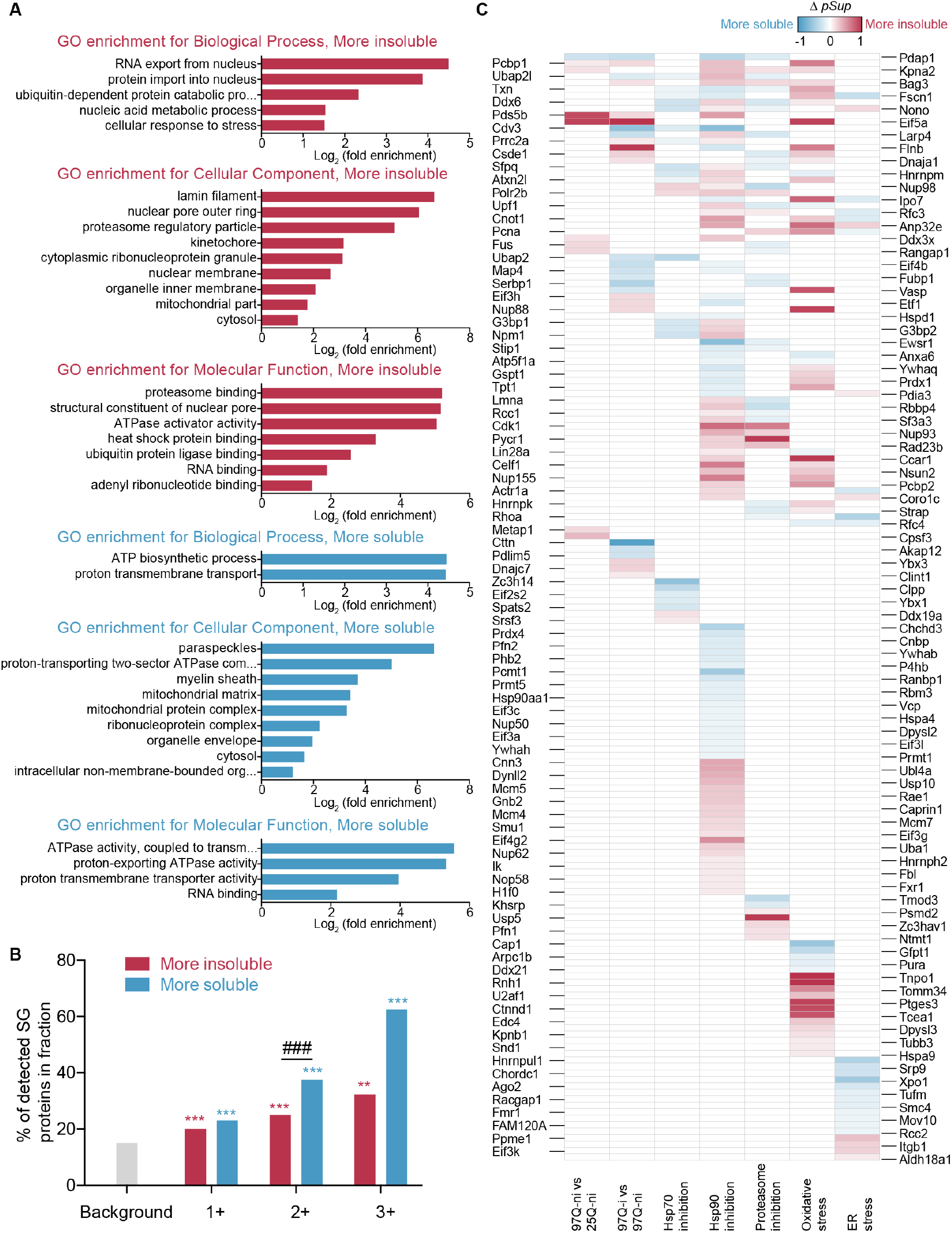
Biological pathways involved in the functional remodeling of proteome solubility. (**A**) GO terms for proteins significantly changed in solubility in two or more stresses. Shown are the top tier GO terms tested with Fisher’s exact test and Bonferroni post hoc testing with a P < 0.05. (**B**) The proportion of detected stress granule proteins using curated list of Markmiller et al. (30) for proteins in indicated number of stresses. (**C**) Barcode graph of stress granule proteins from Markmiller et al. and solubility.

Another striking observation was that 183 of the proteins that become more insoluble in 1 or more of the stresses also became more soluble in 1 or more of the other stresses (**Supplementary Table 7**). This finding suggests that these proteins are functionally tailored to different roles in different stresses. Clues to the key pathways involved came from examination of the proteins that became more insoluble in at least 3 stresses, revealing GO enrichments for Nucleocytoplasmic Transport and Cellular Response to Stress, which are consistent with stress and prior findings that nuclear-cytoplasmic transport is altered in neurodegenerative disease settings (**Fig 4C**) (32, 72–77). Another finding was the profound enrichment of stress granule proteins in both the proteins that become more insoluble, as well as those that become more soluble in at least 3 stresses (**Fig 4C**). Indeed, there was a significant enrichment of proteins involved in stress granules that become both more insoluble and more soluble across one or more stresses (**Fig 5B**). Curiously, a majority of the stress granule proteins displayed differentially altered solubility depending on the stress (**Fig 5C**). These observations indicate a great diversity, dynamism and heterogeneity in the complexes that are formed by stress granule proteins and are suggestive of an elaborate tailoring of the functional responses of the stress granule structures to the stress.

## DISCUSSION

In this study we found that about one-quarter of the proteome is involved in the proteome solubility changes in the mouse neuroblastoma cell in response to an array of stresses to protein homeostasis. Most of the increases in insolubility were counteracted by increases in solubility, as part of a regulatory response to rebalance homeostasis that involved a large functional remodeling of protein-ligand interactions. Our results also suggest that while many proteins involved in protein quality control change as part of the regulatory response, there was almost no common subproteome that appeared to be highly vulnerable to misfolding and aggregation when protein quality control resources are overwhelmed or depleted by different stresses. Specifically, no individual protein consistently became more (or less) soluble under a range of different stresses that have been previously reported to unbalance protein homeostasis. It is therefore apparent that protein aggregation arising from the perturbation of protein quality control under these conditions accounts for only a minor fraction of the changes in the proteome solubility (probably less than 5%). Our results thus reveal an important principle underlying the robustness of protein homeostasis system, namely that there is no bellwether set of proteins that are profoundly metastable, including obvious candidate proteins such as TDP-43 or tau which are commonly mislocalized and-or misfolded in neurodegenerative disease contexts (78, 79). Instead, there is a wider metastable subproteome that underlies specific pathological responses to different stresses. These findings complement those of a recent large-scale proteomic study that identified a small number of endogenous proteins that are prone to aggregation in mouse models of different neurodegenerative diseases, including AD, PD and ALS, indicating the presence of metastable subproteome (80). We also note that our results are compatible with the possibility that aggregation occurs more non-specifically if protein folding is generally perturbed. In this case, small accumulations of each individual protein (e.g. 5%) would most likely be too small to detect with our methodology.

Overall we observed remarkable changes in solubility (both upwards and downwards) involving about one quarter of the proteome for one or more stresses. The signatures for the stresses were generally distinct and are suggestive of widespread remodeling of protein complexes involved in responding to the stress, as evidenced by the GO enrichment in protein quality control-related terms. In this light, we observed the strongest association with stress granule proteins and folding stress as the major activator of quality control systems. Of the proteins not known to be associated with stress granules that become less soluble, this includes chaperone Bag6, proteasome subunit Psde and Vdac2, which mediates Bax translocation in apoptosis (81). Our data point to the robust activation of the quality control system and its remodelling into different sized complexes as a key functional response. In concert, these responses appear to robustly buffer misfolded proteins from actually aggregating. These findings are consistent with a prior study that found heat shock in yeast led to an adaptive autoregulatory assembly and disassembly of protein complexes and minimal aggregation from denatured endogenous proteins (14). Our results extend from this finding to show that such responses are generally applicable to stress and that each stress provides a unique pattern of response.

One of the intriguing findings was the heterogeneity and dynamism of stress granule proteins. There are now at least 238 proteins that have been curated to reside in stress granules (30). It is apparent that there is compositional heterogeneity in the assembly of stress granules (30, 82) and our data further suggests an even more diverse level of heterogeneity and specificity to different forms of stress than currently understood.

The protein seen in 5 stresses, Asns, has been reported to form filaments in yeast under stress (83), suggesting it also is functionally remodeled in response to the stress. When we consider the solubility changes of proteins found in at least 3 stresses in terms of KEGG pathways, we observed a clustering into core metabolic pathways of Lipid Metabolism, Carbohydrate Metabolism, Nucleotide Metabolism, Amino Acid Metabolism and Energy Metabolism (**Fig S6**). The clusters included proteins that increased and decreased solubility and suggests that the remodeling of the proteome solubility is functionally linked to core responses linked to the protein homeostasis stress induction. The key point here is that there is evidence that enzymes involved in metabolism pathways, including specifically those found in our study as hotspots for changes in solubility, form molecular condensates from phase separation in yeast and other cell models (84). This finding suggests a link between metabolic responses, stress granule formation and proteome solubility remodeling involving a quarter of the proteome.

With respect to the metastable subproteome, filamin C (Flnc) stands out as a potential bona fide bellwether protein. Flnc was observed to become more insoluble in 3 stresses (ER stress, oxidative stress and proteasome inhibition). Mutations in Flnc cause myofibrillar myopathies, which are characterized by protein deposits and a defective ubiquitin-proteasome system and oxidative stress (85). Flnc is a cytoskeletal protein that has been shown to aggregate in cells that lack activity of small heat shock protein HspB7 (86). We also observed a binding partner of Flnc, Desm, to be more soluble in four stresses (but of note was not seen in all the stress datasets). This result would be expected if Flnc misfolds and aggregates and is unable to act as an appropriate scaffold to interacting partners. Further evidence is that other genes that cause myofibrillar myopathies when mutated, are Desm, and chaperones DnajB6, HspB8, Cryab and Bag3 which suggest these chaperones are critical to stabilizing the folded state of Flnc (87). We also noted Bag3, which also co-localizes in stress granules, as one of our proteins that become more insoluble in at least 3 stresses. Thus, it remains possible that the machinery involving Bag3 and other stress granules is very competent at buffering against aggregation of misfolded proteins under stress (88), and that Flnc might be one of the first proteins vulnerable to aggregation under prolonged stress.

## AUTHOR CONTRIBUTIONS

X.S. designed and performed the experiments, analyzed the data and helped write the manuscript. S.N. helped perform the proteomics experiments. D.E.V.P., D.B.A, and G.V. performed some of the bioinformatic analysis. M.V. oversaw some of the bioinformatic analysis and preparation of the manuscript. G.E.R. oversaw the proteomics experimental design and analysis. D.M.H. oversaw the project, the design of the experiments, interpretation of the data and drafting of the manuscript. All authors helped draft the manuscript.

## METHODS

### Cell lines

Neuro2a cells, obtained originally from the American Type Culture Collection, were grown in medium consisting of Opti-MEM, 100 U/mL penicillin and 100 μg/mL streptomycin (Life Technologies), 10% (v/v) fetal bovine serum (Thermo Fisher Scientific), 1 mM glutamine in a humidified incubator with 5% atmospheric CO2 at 37 °C.

### DNA vectors and constructs

The constructs with pGW1 vector expressing mCherry as well as Httex1 with 25Q or 97Q were gifts from Steven Finkbeiner, prepared as described previously (89).

### Stress conditions

For the Hsp70 and Hsp90 activity inhibition experiments, Neuro2a cells were cultured in medium supplemented with 20 μM Ver155008 (Sigma) for 18 hours (h) or 800 μM novobiocin (Sigma) for 6 h. To block the proteolytic activity of 26S proteasome complex, cells were stressed in medium supplemented with 5 μM MG132 (Sigma) for 18 h. To induce oxidative stress, cells were treated with 500 μM sodium arsenite (Sigma) for 1 h. To induce ER stress, cells were treated with 5 μM tunicamycin (Sigma) for 18 h. For Httex1 stress, Neuro2a cells were transiently transfected with the vectors using Lipofectamine 2000 reagent (Invitrogen). The cell seeding density was 9 × 10^6^ for each T150 flask. After 24 h, cells were transfected with 120 μL Lipofectamine 2000 and 48 μg vector DNA, as per the manufacturer’s instructions (Invitrogen). The medium was replaced with fresh Opti-MEM medium the following day. After 48 h post-transfection, cells were harvested for flow cytometry sorting.

### Flow cytometry and sorting

Httex1 transfected cells were first rinsed in PBS and harvested by resuspension in PBS with a cell scraper. Cells were pelleted at 120 *g* for 6 min and resuspended in 2 mL PBS supplemented with 10 units/mL DNase I and filtered through 100 μm nylon mesh before analysis by flow cytometry. Cells were analyzed by a FACS ARIA III cell sorter (BD Biosciences) equipped with 405-nm, 488-nm, 561-nm and 640-nm lasers. Side and forward scatter height, width, and area were collected to gate for single live cell population as described previously (24). Data were also collected for pulse height, width, and area of mCherry with a (610/20) filter. The i and ni population was gated by PulSA as described previously (24). To match for expression, cells were further gated to the same median mCherry intensity by varying the window of expression. The Httex1 25Q and 97Q samples were sorted in parallel across four days and used as four matched replicates. For replicate 1, 0.8 million cells were collected for each sample (25Q-ni, 97Q-ni, and 97Q-i); for replicate 2, 1 million cells were collected for each sample; for replicate 3, 1, 0.8 and 0.8 million cells were collected for 25Q-ni, 97Q-ni and 97Q-i, respectively; for replicate 4, 1, 1 and 1.1 million cells were collected for 25Q-ni, 97Q-ni and 97Q-i, respectively. The targeted population was directly sorted into PBS. Cells were snap frozen in liquid nitrogen and stored at − 80 °C for future use.

### Cell fractionation by ultracentrifugation

For all drug treatments (Hsp70, Hsp90, and proteasome inhibition, oxidative stress and ER stress), Neuro2a cells were collected from T75 flasks by first rinsing in 10 mL PBS, followed by resuspension in 10 mL PBS with a cell scraper and gentle pipetting. Cells were pelleted at 120 *g* for 6 min and resuspended in 1 mL ice-cold PBS. After the resuspension mixtures were transferred to chilled 1.5 mL Eppendorf tubes, cells were again pelleted at 120 *g* for 6 min. The resultant pellets were snap frozen in liquid nitrogen and kept at − 80 °C until use. For all stress experiments, each cell pellet was resuspended in 500 μL (200 μL for Httex1 samples) ice-cold Buffer 1 (50 mM Tris-HCl, 150 mM NaCl, 1% (v/v) IGEPAL CA-630, 10 units/mL DNase I, 1:1,000 protease inhibitor) and lysed by forcing through a 27 G needle for 25 times, followed by 31 G for 10 times. After initial lysis, 1 μL (0.8 μL for Httex1 samples) of 0.5 M EDTA was added to each Eppendorf tube to reach a final concentration of 2 mM for EDTA. The cell lysate was then split into two aliquots: a 300 μL (or 120 μL for Httex1 samples) aliquot and the rest left as the representative of the total (T) sample. The 300 μL (or 120 μL for Httex1 samples) aliquot was transferred to a chilled 1.5 mL ultracentrifuge tube and centrifuged at 100,000 *g* for 20 min at 4 °C. 250 μL (or 100 μL for Httex1 samples) of the resulting supernatant was transferred to a new Eppendorf tube and became the supernatant (S) sample. The pellet was washed in 500 μL (or 180 μL for Httex1 samples) Buffer 1, and centrifuged at 100,000 g for 20 min at 4 °C. After ultracentrifugation, the washing buffer was carefully removed without disturbing the pellet. This was repeated for three times. The washed pellet was designated as the pellet (P) fraction. The volume of the T sample was measured by a pipette. The T and S samples were resuspended in the same volume of Buffer 2 (Buffer 1, 4% SDS, 4 mM DTT (1,4-dithiothreitol)) as their initial volume (measured for T, 250 μL (or 100 μL for Httex1) for S), followed by boiling for 20 min at 95 °C. The P fraction was first resuspended in 50 μL Buffer 2 and transferred to an Eppendorf tube, followed by another 50 μL Buffer 2 to wash the wall of ultracentrifugation tube and then added to the Eppendorf tube. The pellet was then boiled for 20 min at 95 °C. The concentration of proteins from T, S, and P fractions was measured by BCA assay by a dilution of 1:20 in PBS according to the manufacturer’s instruction (Thermo Fisher Scientific).

### Protein sample preparation for mass spectrometry

100 μL (approximately 100 μg) of proteins from each sample was treated with 10 mM 1,4-dithiothreitol (DTT) for 1 h to reduce thiols, followed by alkylation with 55 mM iodoacetamide (IAA) for 1 h at 37 °C. Proteins were then extracted by chloroform/methanol precipitation method (90). Briefly, 100 μL chloroform was added to each sample and vortexed, followed by the addition of 300 μL H_2_O and vigorous vortexing. The mixture was then spun at 21,000 *g* for 1 min. The upper aqueous layer was carefully removed without disturbing the precipitated protein at the interface. Subsequently, 300 μL methanol was added and vortexed, followed by centrifugation for 1 min at 21,000 *g*. The supernatant was removed down to a drop so as not to disturb the pellet. The resultant pellet was air dried and dissolved in 100 μL 8 M urea, 50 mM triethylammonium bicarbonate (TEAB) by vigorous vortexing. The solution was then adjusted to a final concentration of 1 M Urea, 100 mM TEAB with addition of 700 μL 100 mM TEAB solution, followed by digestion with trypsin at 1:40 (enzyme: protein ratio) and incubation overnight at 37 °C. The resultant peptides were desalted with a solid-phase extraction (SPE) procedure. The peptide solution was acidified to have 1% (v/v) formic acid. The cartridge (Oasis HLB 1 cc Vac Cartridge, 10 mg sorbent, Waters Corp., USA) was pre-washed with 1 mL of 80% acetonitrile (ACN) containing 0.1% trifluoroacetic acid (TFA) and washed twice with 1.2 mL of 0.1% TFA. The acidified peptides were loaded on the cartridge, followed by washing with 1.5 mL of 0.1% TFA twice. The sample was eluted with 800 μL of 80% ACN containing 0.1% TFA and collected in a fresh 1.5 mL Eppendorf tube, followed by lyophilization by freeze drying (Virtis, SP Scientific). Freeze dried peptides were resuspended in 100 μL of distilled water and quantified by BCA assay (Thermo Fisher Scientific) according to the manufacturer’s instruction. 25 μL of each sample was then diluted into a final volume of 100 μL containing 100 mM TEAB, prior to reductive dimethyl labelling. 4 μL of 4% (v/v) formaldehyde – CH_2_O (light label), CD_2_O (medium label), and/or C^13^D_2_O (heavy label) were added, followed by addition of 4 μL of 0.6 M sodium cyanoborohydride and incubation for 1 h at 20°C. The reaction was then quenched by addition of 16 μL of 1% ammonium hydroxide followed by 8 μL of neat formic acid.

### NanoESI-LC-MS/MS analysis

Peptides were analyzed by liquid chromatography-nano electrospray ionization- - tandem mass spectrometry (LC-nESI-MS/MS) using either an Orbitrap Fusion Lumos mass spectrometer (Thermo Fisher Scientific, San Jose, CA) or Orbitrap Q Exactive Plus mass spectrometer (Thermo Fisher Scientific) Bremen, Germany. The nano-LC system, Ultimate 3000 RSLC (Thermo Fisher Scientific, San Jose, CA) was equipped with an Acclaim Pepmap nano-trap enrichment column (C18, 100 Å, 75 μm × 2 cm, Thermo Fisher Scientific, San Jose, CA) and an Acclaim Pepmap RSLC analytical column (C18, 100 Å, 75 μm × 50 cm, Thermo Fisher Scientific, San Jose, CA) maintained at a temperature of 50 °C. Typically for each experiment, 1 μg of the peptide mixture was loaded onto the enrichment column at an isocratic flow of 5 μL min^-1^ of 3% CH3CN containing 0.05% TFA for 5 min before the enrichment column was switched in-line with the analytical column. The eluents used for the LC were water with 0.1% v/v formic acid and 5% v/v dimethyl sulfoxide (DMSO) for solvent A and ACN with 0.1% v/v formic acid and 5% DMSO for solvent B.

For Hsp70 inhibition, proteasome inhibition, and arsenite stress samples, the gradient used (300 nL/min) was from 3% B to 23% B for 89 min, 23% B to 40% B in 10 min, 40% B to 80% B in 5 min and maintained at 80% B for the final 5 min before equilibration for 8 min at 3% B. The gradient for Httex1 samples was from 3% B to 22% B for 102 min, 22% B to 32% B in 15 min, 32% B to 95% B in 10 min and maintained at 95% B for the final 10 min before equilibration for 10 min at 3% B prior to the next analysis. The experiments were performed in positive ionization mode on the Orbitrap Fusion Lumos mass spectrometer. The spray voltages, capillary temperature and S-lens RF level were set to 1.9 kV, 275 °C and 30%. The mass spectrometry data was acquired with a 3-second cycle time for one full scan MS spectra and as many data dependent HCD-MS/MS spectra as possible. Full scan MS spectra were acquired using an m/z range from 400 to 1500 (m/z 375 to 1500 for Httex1 samples), a resolution of 120,000 at m/z 200 using Orbitrap mass analysis, an auto gain control (AGC) target value of 4e5, and a maximum ion trapping time of 50 milliseconds. Data dependent HCD-MS/MS of peptide ions (charge states from 2 to 5) was performed using an m/z isolation window of 1.6, an AGC target value of 5e4 (1e4 for Httex1 samples), a normalized collision energy (NCE) of 35%, a resolution of 7,500 at m/z 200 with a maximum ion trapping time of 22 milliseconds (proteasome inhibition), or a resolution of 15,000 at m/z 200 with a maximum ion trapping time of 60 milliseconds (Hsp70 inhibition and arsenite stress) using Orbitrap mass analysis, or a maximum ion trapping time of 60 milliseconds and a ion trap mass analyzer at rapid scan rate with a q value of 0.25 (Httex1 samples). Dynamic exclusion was used for 30 s.

For Hsp90 inhibition and ER stress samples, the gradient used (300 nL/min) was from 3% B to 23% B for 89 min, 23% B to 40% B in 10 min, 40% B to 80% B in 5 min and maintained at 80% B for the final 5 min before equilibration for 8 min at 3% B prior to the next analysis. The MS experiments were performed in positive ionization mode on the Orbitrap Q Exactive Plus mass spectrometer. The spray voltages, capillary temperature and S-lens RF level were set to 1.9 kV, 250 °C and 50%. Full scan MS spectra were acquired using an m/z range from 375 to 1400, a resolution of 70,000 (Hsp90 inhibition) or 120,000 (ER stress) at m/z 200, an AGC target value of 3e6, and a maximum ion trapping time of 50 milliseconds. The top 15 data dependent HCD-MS/MS of peptide ions (charge states from 2 to 5) was performed using an m/z isolation window of 1.2 (Hsp90 inhibition) or 0.7 (ER stress), m/z scan range of 200 – 2000, a AGC target value of 1e5, a stepped NCE of 28%, 30% and 32% (Hsp90 inhibition) or a NCE of 30% (ER stress), a resolution of 35,000 (Hsp90 inhibition) or 30,000 (ER stress) at m/z 200 with a maximum ion trapping time of 120 milliseconds. All mass spectrometry data were acquired using the Orbitrap mass analyzer. Dynamic exclusion was used for 30 s.

Data analysis was performed using Proteome Discoverer (version 2.1.0.81; Thermo Scientific) with the Mascot search engine (Matrix Science version 2.4.1). Database searches were conducted against the Swissprot Mus musculus database (version 2015_07: Jun-24, 2015; 548872 entries), with 20 ppm MS tolerance, 0.2 Da MS/MS tolerance and 2 missed cleavages allowed. Variable modifications were used for all drug treatment experiments: oxidation (M), acetylation (Protein N-term), dimethylation (K), dimethylation (N-Term), 2H(4) dimethylation: (K), 2H(4) dimethylation (N-term); for Httex1 samples, the modifications of 2H(6)13C(2) dimethylation (K), 2H(6)13C(2) dimethylation (N-term) were also used. A fixed modification used for all experiments was carbamidomethyl (C). The false discovery rate (FDR) maximum was set to 0.5 % at the peptide identification level and 1 % at the protein identification level. Proteins were filtered for those containing at least two unique peptides in all n=4 biological replicates. Peptide quantitation was performed in Proteome Discoverer v.2.1.0.81 using the precursor ion quantifier node. Dimethyl labelled peptide pairs were established with a 2 ppm mass precision and a signal to noise threshold of 3. The retention time tolerance of isotope pattern multiplex was set to 0.8 min. Two single peak or missing channels were allowed for peptide identification. The protein abundance in each biological replicate was calculated by the mean of the top 3 unique peptide abundances that were used for quantitation. For total and pellet proteome analysis, the median protein ratio (i.e., stress/control) of each replicate was normalized to 1; for supernatant proteome, the median protein ratio (i.e., supernatant/total) of each replicate was normalized to 0.8 based on the proportion (~ 0.8) of the supernatant fraction out of the total lysate determined by the BCA assay. One-sample t-test (for ratios generated from the total and pellet proteome) and two-sample t-test (for *pSup* values generated from the supernatant proteome) was performed to calculate the statistical significance with Perseus.

### Bioinformatics analysis and visualization using Cytoscape

For GO analysis, data was analyzed by the PANTHER overrepresentation test (PANTHER version 14.0, 20181113) using the PANTHER GO-Complete Biological Process, Cellular Component and Molecular Function dataset. Fisher’s exact test using the Bonferroni correction for multiple testing was performed. Results were filtered for p < 0.05.

For protein interaction network analysis, Cytoscape 3.7.1 (91) was used. Excel files containing the input data were uploaded to Cytoscape. Protein interaction network was generated with built-in String (v11.0) (92) using active interaction sources parameters on for Experiments, Databases, Co-expression neighborhood, Gene Fusion and Cooccurrence. The minimum required interaction score setting was 0.4 (medium confidence). Total protein abundance ratios or protein solubility changes (including Δ *pSups* and log_2_(pellet ratios) with maximum and minimum values normalized to 1 and - 1) were used as node color attributes. Nodes were manually re-arranged based on GO terms. Adobe illustrator was used to annotate GO terms on the protein interaction network.

The cut-offs were selected to reflect the best separation identified between classes, for each stress. They were originally evaluated for their signal to noise/information using Random Forest under leave-one-out cross-validation. The values are 0.15 for Δ *pSup* and 0.5849 for log2(pellet ratio) with p < 0.05 for all stresses with exceptions for proteasome inhibition (0.1 and 0.8074) and oxidative stress (0.05 and 0.5849).

Venn diagrams of protein hits across different treatments were performed using Venny (http://bioinfogp.cnb.csic.es/tools/venny/). Heat map and pairwise comparison of proteins hits between different treatments was performed using Heatmapper (http://www1.heatmapper.ca/pairwise/).

Protein identifiers (UNIPROT ids) were used to map selected proteins (more soluble/insoluble sets) to metabolic pathways and create metabolic hot spot maps using KEGG mapper (https://www.genome.jp/kegg/tool/map_pathway2.html).

### Statistical analysis

The details of the tests were indicated in the figure legends. Significant results were defined for *p* < 0.05.

### Data availability

The mass spectrometry proteomics data have been deposited to the ProteomeXchange Consortium via the PRIDE [1] partner repository with the dataset identifier PXD014420.

## LIST OF FIGURES AND SUPPLEMENTAL DATA

Supplementary Data 1. Gene Ontology terms associated with the abundance and solubility datasets.

Supplementary Table 1. Httex1 proteomics dataset

Supplementary Table 2. Hsp70 inhibition proteomics dataset

Supplementary Table 3. Hsp90 inhibition proteomics dataset

Supplementary Table 4. Proteasome inhibition proteomics dataset

Supplementary Table 5. Oxidative stress proteomics dataset

Supplementary Table 6. ER stress proteomics dataset

Supplementary Table 7. Commonalities of proteins that are significantly altered in solubility between stresses

### Supplementary Note 1. Expanded discussion of proteins that change in expression level by mutant Httex1 aggregation state

For cells with mutant 97Q Htt but lacking inclusions (97Q ni), we observed Lgmn, which is an endopeptidase responsible for cleaving full length Htt into toxic N-terminal Htt fragments (93, 94) and Msh2 which upon deletion suppressed somatic Htt CAG expansion mutations and aggregation of Htt in HdhQ111 mice (95). We also saw proteins that mediate apoptosis downregulated including Plk1 (96–98) and its activator Ociad1is (99). Also downregulated was mitophagy mediator Tbc1d15, which is a mitochondrial RAB-GTPase activating protein (31).

Up-regulated proteins included mitochondrial import protein Timm13 (100), which may indicate activation of a recently described mitochondrial quality control mechanism to clear misfolded protein (101).

For proteins that are altered upon 97Q aggregation, most of the handful of proteins were upregulated and many had reported roles in degradation. Interestingly, the proteins upregulated suggested a role in stabilizing, rather than degrading Httex1. This included deubiquitinating enzyme Usp19, which has been previously reported to stabilize mutant Httex1 levels and promote its aggregation into less toxic aggregates by Hsp90 (102, 103). Rock2 was also increased, which has previously been suggested to slow degradation of Httex1 (104). Rabgap1 was also seen, which is an activator of mTORC1 that has many cellular functions including inhibiting autophagy. Of note is its relevance to Huntington pathology where TORC1 hyper-activation has been reported (105).

**Supplementary Fig 1. (Relates to Fig 2).**
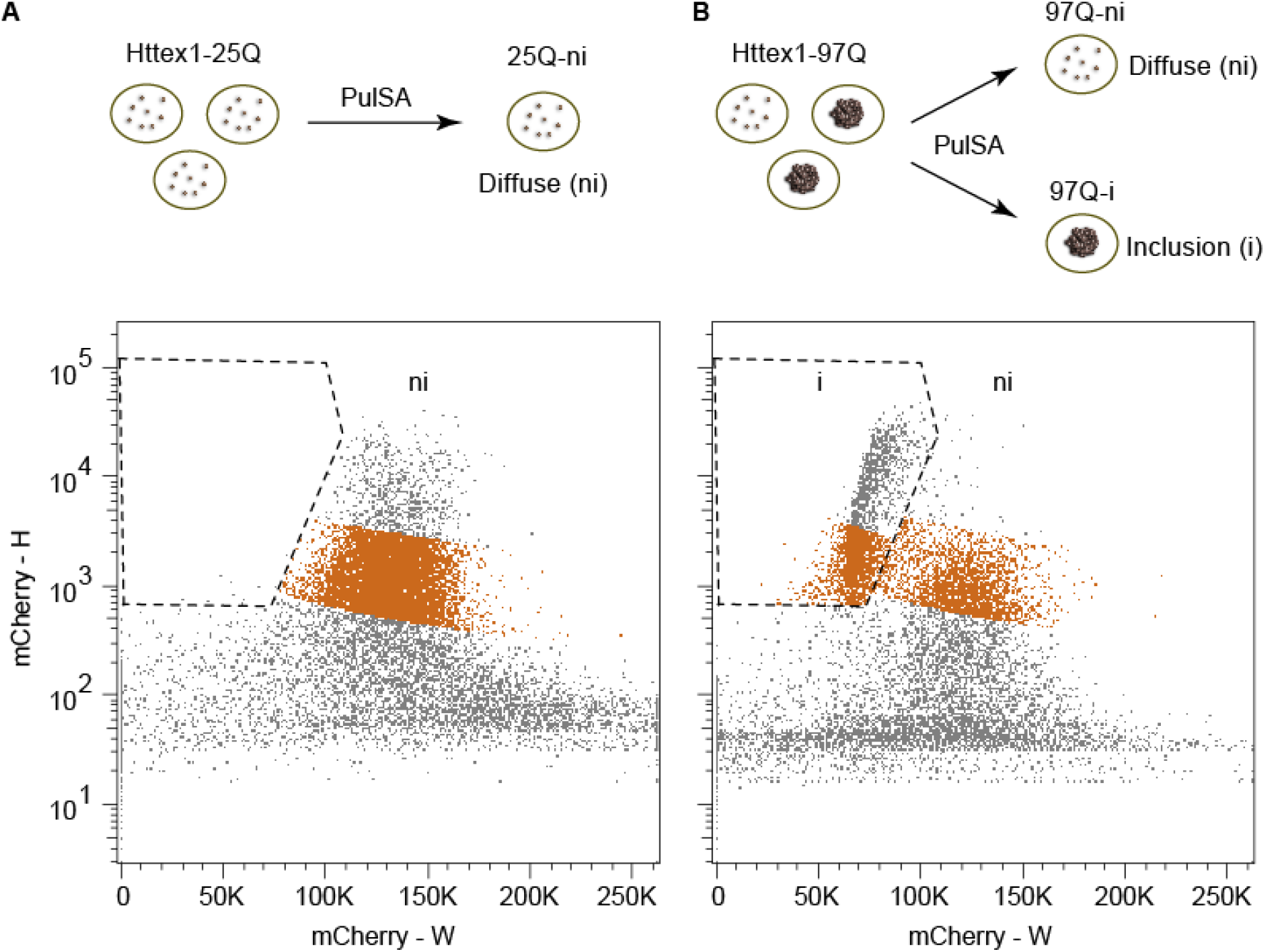
Strategy to sort cells with Httex1-mCherry inclusions from those without inclusions by PulSA. Schematic and flow cytometry gates for Httex1(25Q) (**A**) and Httex1(97Q) (**B**) fusions to mCherry. PulSA was applied on the mCherry Width and Height fluorescence parameters. Cells were selected to maintain matched median fluorescence values within the i gate and ni gates (orange-brown coloured data).

**Supplementary Fig 2. (Relates to Fig 3).**
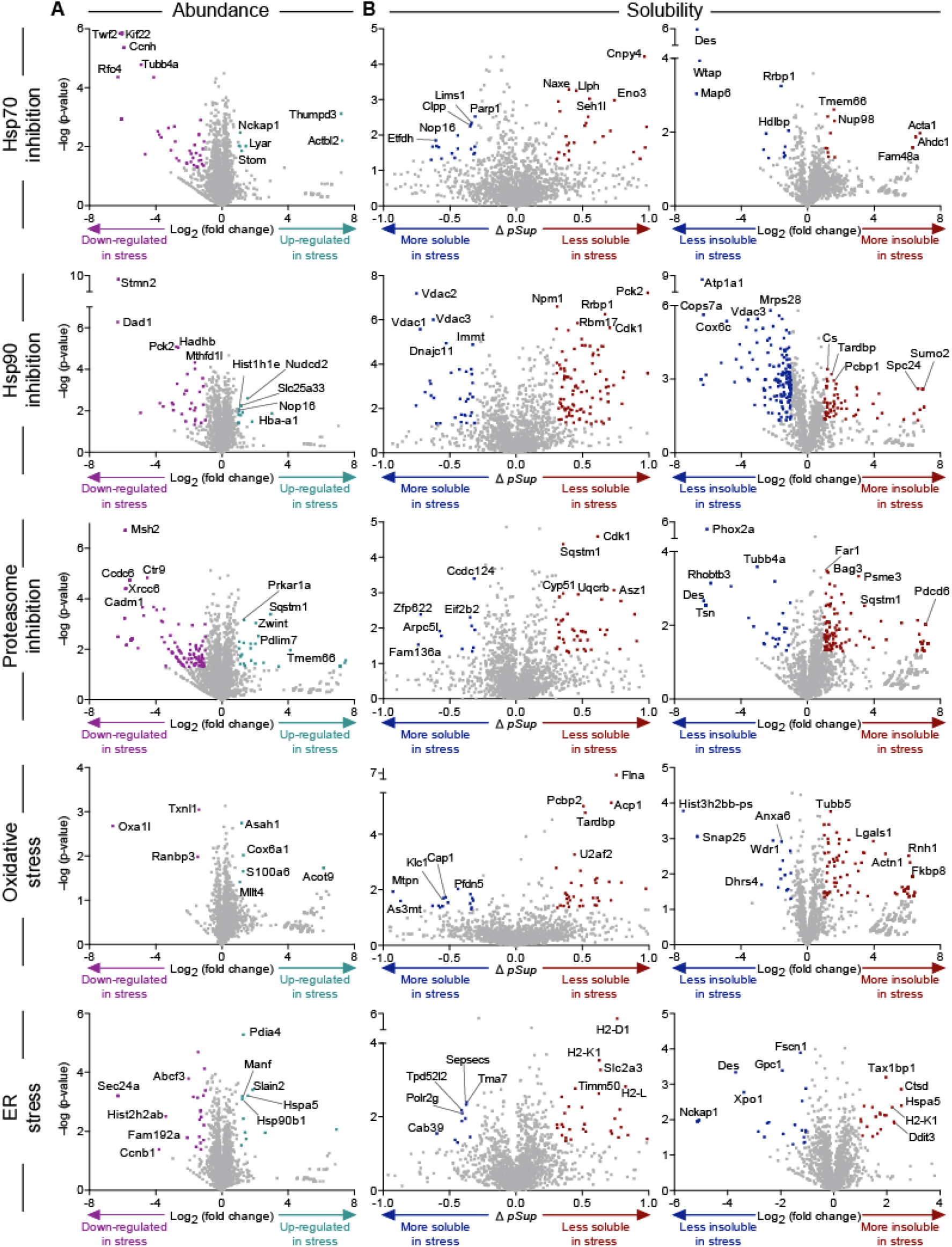
Proteome abundance and solubility changes arising from protein homeostasis stresses. Volcano plots of protein abundances and solubility for stresses versus controls including hsp70 with ver-155008, inhibition of hsp90 with novobiocin, inhibition of the proteasome with MG132, oxidative stress with sodium arsenite, and ER stress with tunicamycin. Grey-colored points indicate proteins below the threshold of significance (2-fold change and t-test P value of 0.05 for abundance and *Δ pSup* value of 0.3, Pellet ratio log_2_ value of 1 and t-test P value of 0.05). 5 most changed proteins in each direction are shown.

**Supplementary Fig 3. (Relates to Fig 3).**
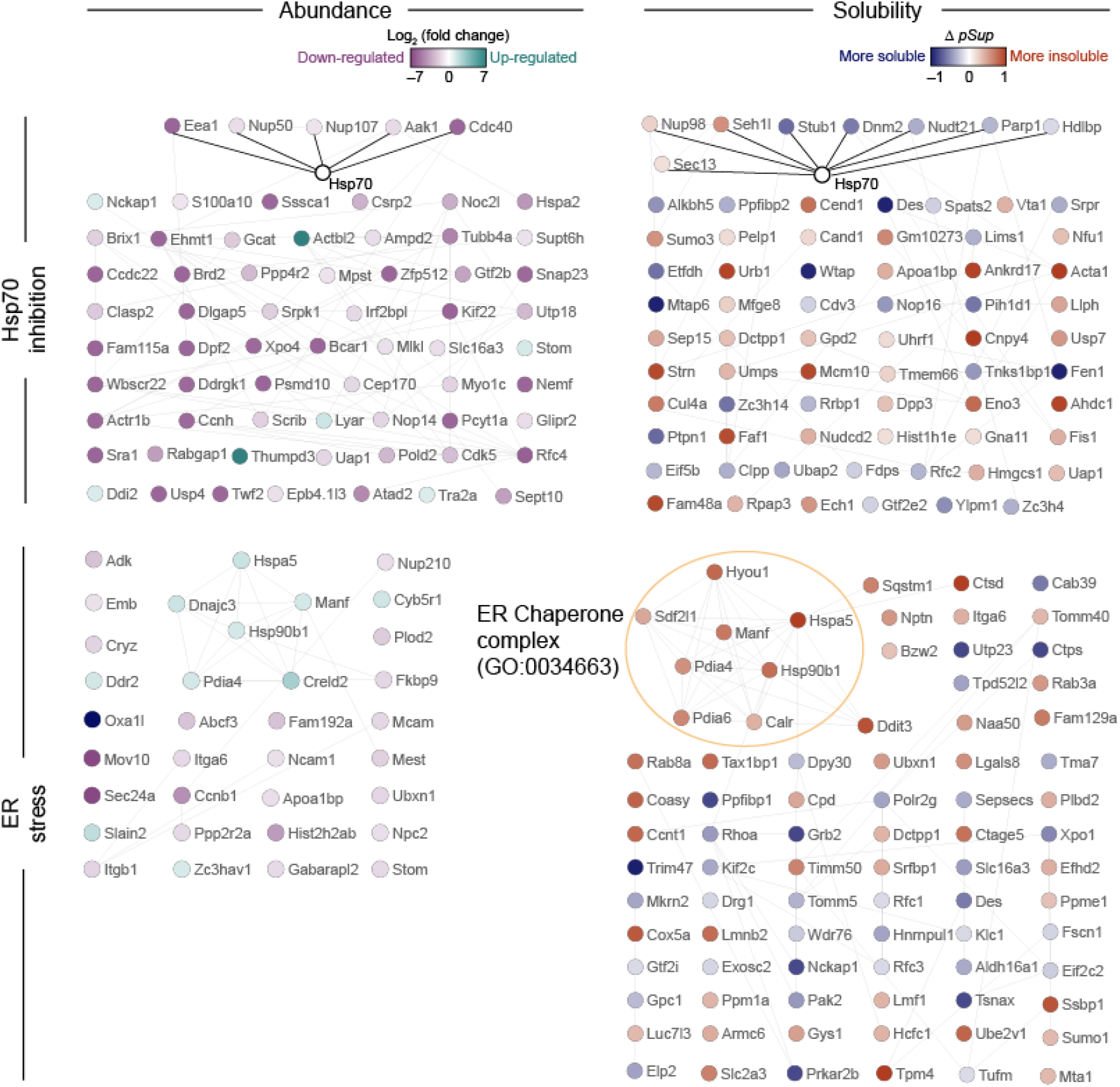
Impact of Hsp70 inhibition and ER stress on proteome abundance and solubility. Protein:protein interaction analysis of proteins with significant changes in abundance (left panels) or solubility (right panels) due to indicated protein homeostasis stresses was performed built-in String (v.11) in Cytoscape (v3.7.1) at the medium confidence. Selected significantly enriched Gene Ontology terms are annotated. All enriched Gene Ontology terms are included in Supplementary Table S2. For Hsp70 inhibition, Hsp70 interactors are shown as solid black lines.

**Supplementary Fig 4. (Relates to Fig 3).**
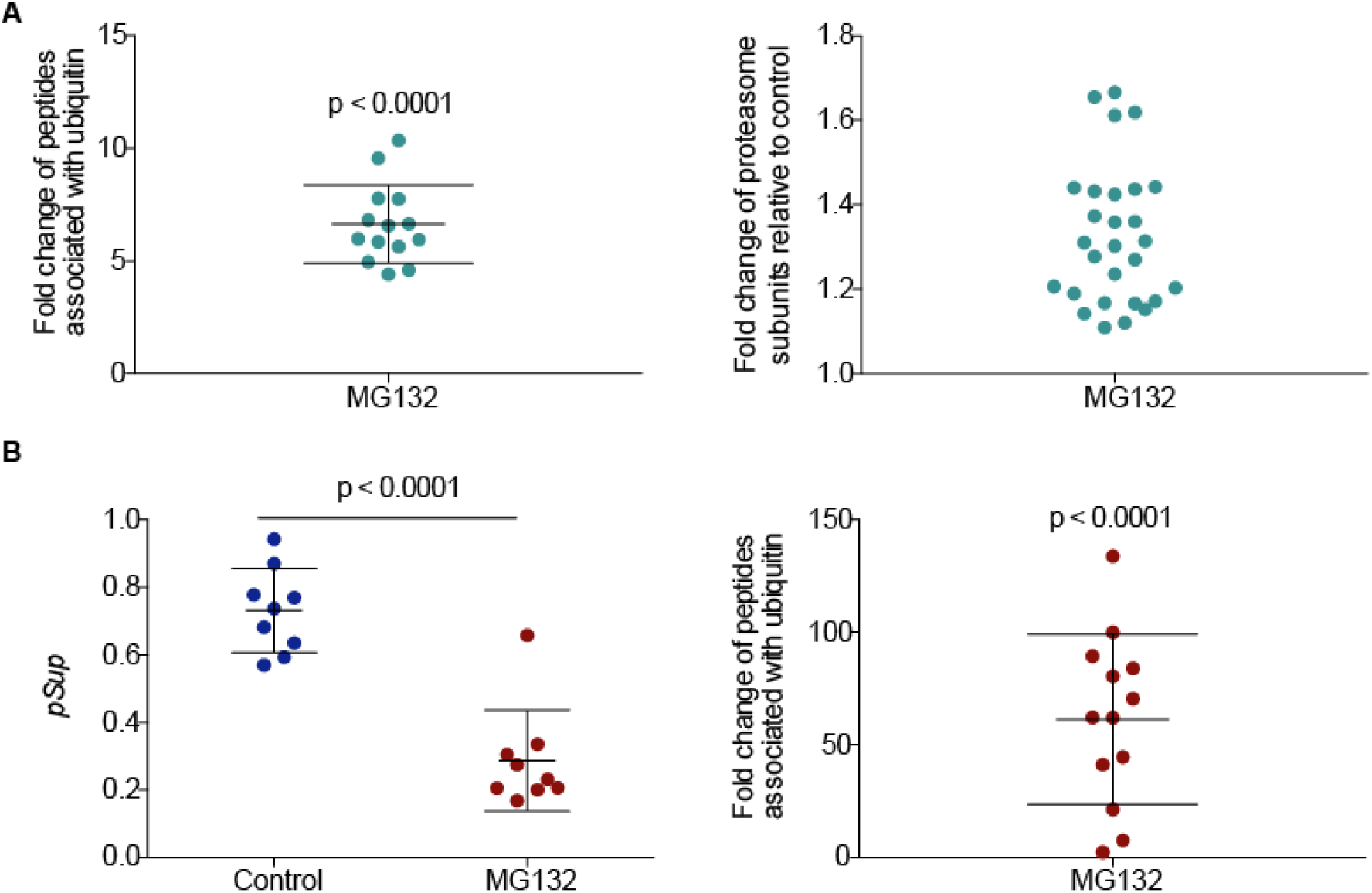
Ubiquitin was up-regulated and had reduced solubility upon proteasome inhibition. (**A**) Left panel shows the relative abundance of peptides generated from ubiquitin after MG132 treatment compared with control. Data are presented as mean +/− SD. Statistical significance was calculated by t-test. Right panel shows the relative abundance of proteasome subunits upon MG132 treatment compared with control. (**B**) Data shows the solubility changes of ubiquitin due to MG132 treatment with (*pSup*) on the left graph and pellet ratio on the right graph. Each data point indicates the average value from four biological replicates for each peptide. Mean +/− SD is shown. P value indicates result of two-tailed t-test.

**Supplementary Fig 5. (Relates to Fig 4).**
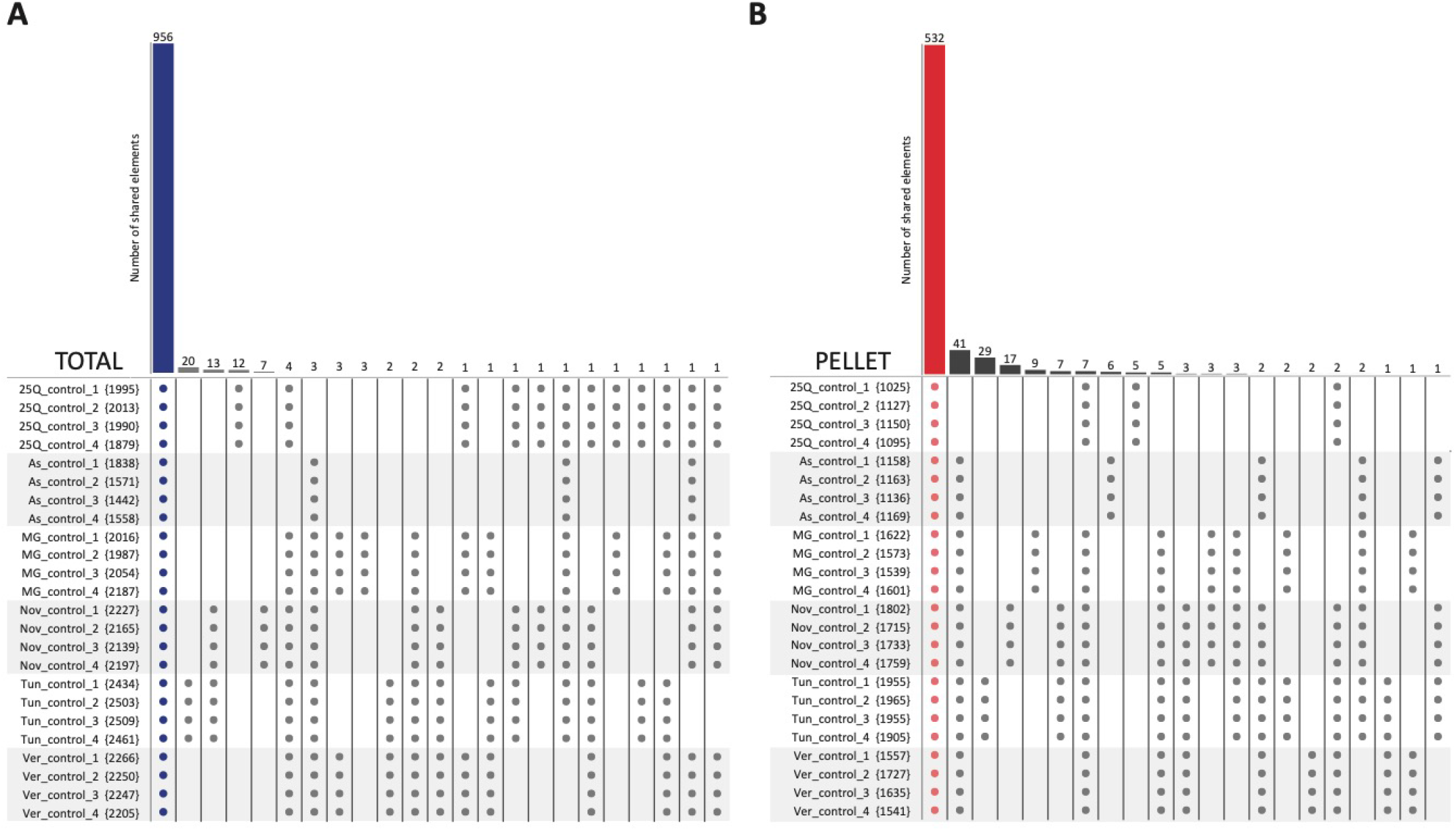
Metastable subroteome under control conditions. Alternative representation of Venn diagrams showing how many proteins are identified consistently across the conditions in the control samples, where consistently implies that the proteins are quantified in 4 out of 4 replicas (dots in the table). The columns of the tables represent the various sets overlaps, reported as the row of the table, where dot means presence of the set and nothing means absence of the set in that overlap. **(A)** Proteins quantified in the total fraction of the control samples for the 6 conditions. The first column (956 proteins, blue bar) shows the number of proteins found in the total fraction for all replicas of all control conditions. All other overlaps are nearly 2 orders of magnitude smaller, hence negligible in comparison. **(B)** Proteins quantified in the pellet fraction of the control samples for the 6 conditions. The first column (532 proteins, red bar) shows the number of proteins found in the pellet fraction for all replicas of all control conditions. All other overlaps are 1 order of magnitude smaller, hence negligible in comparison.

**Supplementary Fig 6. (Relates to Fig 5).**
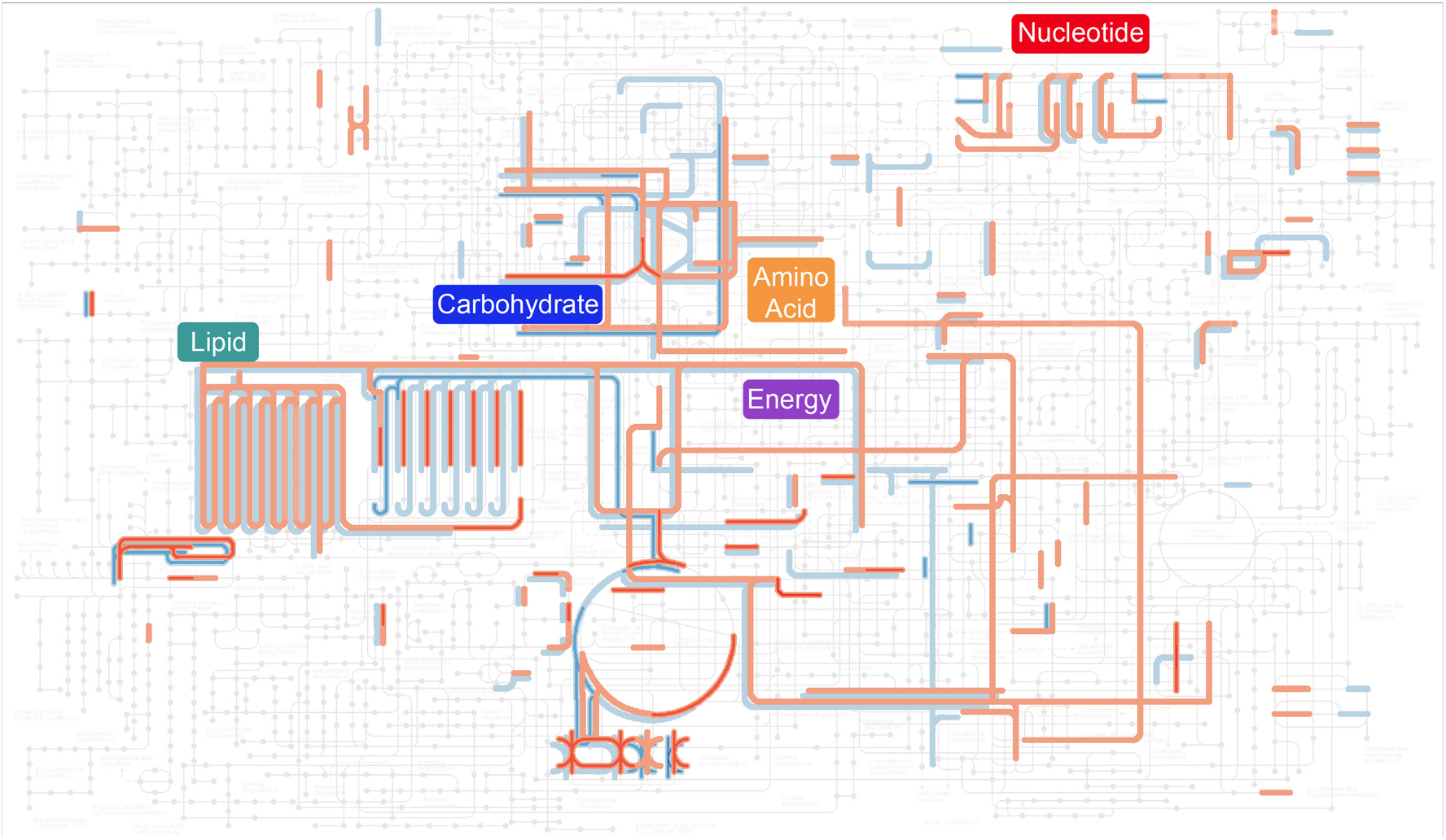
Metabolism hotspots in proteome solubility remodeling. Shown is the KEGG metabolism atlas and colour-coded are pathways with proteins that change solubility in 2 or more stresses. Red indicates more insoluble protein enrichment and blue more soluble enrichment.

## Footnotes

1 This includes the sum of proteins seen in the 25Q-ni vs 97Q-ni and 97Q-ni vs 97Qi treatments

2 Note we were unable to measure solubility for the Huntington’s Disease cell model due to small yields of cells after flow cytometry sorting.

